# Discovery of two novel *Torque Teno viruses* in *Callithrix penicillata* provides insights on *Anelloviridae* diversification dynamics

**DOI:** 10.1101/2022.06.30.498111

**Authors:** Matheus Augusto Calvano Cosentino, Mirela D’arc, Filipe Romero Rebello Moreira, Liliane Tavares de Faria Cavalcante, Ricardo Mouta, Amanda Coimbra, Francine Bittencourt Schiffler, Thamiris dos Santos Miranda, Gabriel Medeiros Viana, Cecilia A Dias, Antonizete R Souza, Maria Clotilde Henriques Tavares, Amilcar Tanuri, Marcelo Alves Soares, André Felipe Andrade dos Santos

## Abstract

The development of high throughput sequencing (HTS) technologies and metagenomics protocols deeply impacted the discovery of viral diversity. Moreover, the comprehension of evolution and immunology of the Neotropical primates (NP) and their susceptibility to infectious diseases are central for the characterization of the scope of etiological agents that may impact global health, due to their evolutionary proximity to Old World primates, including humans. In the present work, novel anelloviruses were detected and characterized through HTS protocols in the NP Callithrix penicillata, the common black-tufted marmoset. De novo assembly of generated sequences was carried out and a total of 15 contigs were identified with complete Anelloviridae ORF1 gene, two of them including a flanking GC-rich region, confirming the presence of two whole novel genomes of ~3kb. The identified viruses were monophyletic within the Epsilontorquevirus genus, a lineage harboring previously reported anelloviruses infecting hosts from the Cebidae family. The genetic divergence found in the new viruses characterized two novel species, named *Epsilontorquevirus callithrichensis I* and *II*. The phylogenetic pattern inferred for the *Epsilontorquevirus* genus was consistent with the topology of their host species tree, echoing a virus-host diversification model observed in other viral groups. This study expands the host span of *Anelloviridae* and provides insights into their diversification dynamics, highlighting the importance of sampling animal viral genomes to obtain a clearer depiction of their long-term evolutionary processes.

## 2. Introduction

The development of high throughput sequencing (HTS) technologies and metagenomics protocols deeply impacted the discovery of viral diversity ^1,2^. Once a matter of contention, the establishment of novel viral species and higher taxonomic groups based solely on sequencing data became commonplace. Prospective studies describing multiple previously unknown viral groups are increasingly common ^3–6^, as are studies expanding the diversity of well-established viral families ^7–10^. While these efforts have broad implications on the comprehension of virus diversity, ecology, evolution and epidemiology, large fractions of the viruses infecting non-model organisms remain to be discovered ^11^. In this regard, Neotropical primates (NP) (infraorder Platyrrhini) stand out as a group with great ecological importance, whose viral diversity hasn’t been fully explored. Only 18% (4/22) and 3.4% (6/178) of NP genera and species, respectively, have been previously assessed through viral metagenomics ^10,12–16^.

Currently, NP comprehend 178 species and 22 genera that occupy a myriad of ecological niches in Central and South Americas^17^. As it is the sister group to Old World primates (OWP) (infraorder Catarrhini), including humans, the comprehension of their evolution, immunology and susceptibility to infectious diseases is central to the characterization of the scope of etiological agents that may impact global health^18^. In this sense, previous investigations have shown that multiple viral families might be found in the virome of NP: novel papillomaviruses were found infecting *Callithrix penicillata*^16^ and *Alouatta guariba*^15^, anellovirus and genomovirus in *Sapajus nigritus* ^10^, smacovirus in *Alouatta caraya* ^12^ and foamy virus in *Brachyteles arachnoides* ^14^ and *Sapajus xanthosternos* ^13^.

The *Anelloviridae* family is one of the most frequently found in metagenomic surveys of NP. This family is composed of non-enveloped virions with circular single-stranded negative sense DNA (ssDNA), whose genomes range from 1.6 to 3.9kb. Currently, the group is composed of 30 classified genera by the International Committee of Virus Taxonomy ^19^. The *Torque Teno Virus* (TTV) was first identified in the 1990s in a post-transfusion hepatitis case of unknown etiology in humans ^20^ and was posteriorly discovered in healthy human populations with prevalence ranging from 10 to 80%^21^. No pathology has been confirmed by its infection, although an association with pulmonary and liver diseases has been suggested^22^. Furthermore, the *Anelloviridae* family is proposed to be a common part of the human virome, with major importance to the immune system maturation^23^. A high prevalence in healthy populations and the lack of clinical signs have also been shown in free-living and captive populations of chimpanzees and gorillas ^24–26^. Although new anelloviruses species have been discovered in NP - such as in *Saguinus oedipus, Aotus trivirgatus*^27^ and *Sapajus nigritus*^10^ - an extensive knowledge gap for anelloviruses infecting NP still remains.

While the diversity and host span of *Anelloviridae* is constantly being updated^9,10,25,27–33^, little is known about the factors driving their diversification dynamics. Specifically, whether the generation of novel viral species is driven by zoonotic spillovers to new host species or co-divergence with long-term host lineages has not been formally assessed. This problem has been addressed by phylogenetic methods for other groups in large scale meta-analysis^34^ as well as in studies focusing on specific viral families^7,15,35–37^, revealing that the mechanisms that generate viral diversity are often determined by viruses genomic architecture and variability. As the discovery of novel viruses allows a broader evaluation of these mechanisms, metagenomic surveys on non-model organisms have allowed the clarification of the evolutionary dynamics of diverse viral families ^13,14,38,39^.

In this study, we characterized the virome of captive *Callithrix penicillata*, the black-tufted marmoset, by HTS and discovered two novel species of anelloviruses, expanding the scope of primate hosts infected by these viruses. Phylogenetic analysis of the novel sequences suggests anellovirus evolution might be driven by the within-species diversity followed by co-divergence model, as suggested for other viral groups ^16,40–42^. Overall, our results shed light on previously unexplored anellovirus diversity and address a hypothesis regarding their diversification dynamics.

## 3. Material and Methods

### 3.1. Sample Collection

The data analyzed herein was generated on a larger virome project using samples from NP housed at the *Centro de Primatologia* of *Universidade de Brasília* (CP/UnB; Brasília - Brazil). Specific viral family data is being published as analyses are finalized and general information can be found in the first publication of the larger project^16^, which characterized papillomavirus in primates from this center. For this study, the data analyzed were sequenced from anal swab samples from sixteen black-tufted marmosets (*C. penicillata*), immersed in phosphate-buffered saline (PBS) and stored at −20°C (Table 1). Samples were collected following the national guidelines and provisions of National Council for Animal Experimentation Control (CONCEA), which included animal welfare standard operating procedures. The procedure was conducted under permission of the Brazilian Institute of Environment and Renewable Natural Resources (IBAMA) under license number 1/53/1999/000006-2 and the project was approved by Ethics Committee on the Use of Animals of Universidade Federal do Rio de Janeiro (reference number 037/14).

**Table 1:** General information of all samples from Callithrix penicillata housed at Centro de Primatologia of Universidade de Brasília (CP/UnB).

| Specie | CP/UnB name | Lab ID | Library Pool ID | Sample Type | Gender | Sexual Maturity | Enclosure |
| --- | --- | --- | --- | --- | --- | --- | --- |
| *Callithrix penicillata* | Pamonha | SWA1 | P1 | Anal Swab | F | Adult | 1 |
| *Callithrix penicillata* | Novela | SWA2 | P1 | Anal Swab | F | Adult | 2 |
| *Callithrix penicillata* | Miuxa | SWA3 | P1 | Anal Swab | F | Adult | 3 |
| *Callithrix penicillata* | Miúda | SWA4 | P1 | Anal Swab | F | Adult | 3 |
| *Callithrix penicillata* | Grelina | SWA5 | P1 | Anal Swab | F | Adult | 4 |
| *Callithrix penicillata* | Rubi | SWA6 | P2 | Anal Swab | F | Adult | 6 |
| *Callithrix penicillata* | Gonia | SWA7 | P2 | Anal Swab | F | Adult | 8 |
| *Callithrix penicillata* | Valina | SWA8 | P2 | Anal Swab | F | Adult | 9 |
| *Callithrix penicillata* | Gora | SWA9 | P2 | Anal Swab | F | Adult | 10 |
| *Callithrix penicillata* | Miuka | SWA10 | P2 | Anal Swab | F | Adult | 10 |
| *Callithrix penicillata* | Gontijo | SWA45 | P7 | Anal Swab | M | Adult | 5 |
| *Callithrix penicillata* | Valente | SWA46 | P7 | Anal Swab | M | Adult | 7 |
| *Callithrix penicillata* | Supimpa | SWA47 | P7 | Anal Swab | M | Adult | 8 |
| *Callithrix penicillata* | Miro | SWA48 | P7 | Anal Swab | M | Adult | 9 |
| *Callithrix penicillata* | Gonero | SWA51 | P7 | Anal Swab | M | Adult | 14 |
| *Callithrix penicillata* | Birrento | SWA53 | P7 | Anal Swab | M | Adult | 16 |

### 3.2. High-Throughput Sequencing

The samples were shipped to the *Laboratório de Diversidade e Doenças Virais* of *Universidade Federal do Rio de Janeiro* (LDDV/UFRJ; Rio de Janeiro - Brazil) and the homogenized PBS aliquots were pooled into three arbitrarily sampled groups (P1, P2 and P7) to maximize sequencing efficiency with 4 to 6 samples each (Table 1). Pooled samples were filtered using Millex-HV 0.45 μm SLHV0135L (Merck, German) and sacarose gradient by ultracentrifugation at 35,000 g, 4°C for 90 minutes was conducted to separate the enriched viral bottom fraction. Supernatants were discarded and the remaining 0.2 mL were digested with DNases (Promega and Epicentre, USA), RNase A (Ambion, USA) and Benzonase (Sigma, USA). The QIAamp MinElute Virus Spin Kit (QIAGEN, Germany) was used for viral nucleic acid extraction, following the manufacturer’s protocol with slight modifications to improve the recovery of total nucleic acids in accordance to the CDC protocol methodology^43^: i) the “Carrier RNA” was not used in the AL Buffer; ii) the protease was resuspended with AVE Buffer, instead of Protease Resuspension Buffer; iii) the washing step with AW1 was not performed; iv) the final elution was done in 20 μL of ultra-pure water. The material was then subjected to a complementary DNA (cDNA) synthesis using the Superscript III First Strand Synthesis Supermix Kit (Thermo Fisher Scientific, USA). The second strand cDNA synthesis was performed using Klenow fragment 3’-5’ exo (New England Biolabs Inc, USA), following the manufacturer’s instructions. Sequencing libraries were constructed using Nextera XT DNA Library Preparation Kit (Illumina, USA), following the standard protocol. In a MiSeq V2 500-cycle cartridge (Illumina), 4 pM of each library were loaded and massive sequencing was conducted on the Illumina MiSeq platform of the Department of Genetics of UFRJ.

### 3.3. Bioinformatic Analysis

Sequencing data was processed with a custom pipeline. Raw sequencing reads were filtered with Fastp v.0.20.1^44^, removing short reads (<50bp) and low-quality (Phred score < 30) bases. Reads for each sample were then de novo assembled with the Meta-spades software v.3.15.3^45^. Diamond v.2.0.14^46^ was used to taxonomically assign both reads and their respective contigs, with an e-value cut-off of 10^−5^ and parameters ‘--more-sensitive’ and ‘--max-target 1’. A similarity search was performed against the complete non-redundant protein (nr) database from NCBI (as of July 27th, 2021). Krona v.2.7.1^47^ was used to generate interactive visualization plots, allowing the fast identification of viral families of interest. To avoid false positive results due to sequencing index-hoping (incorrect assignment of a viral read to a given sample), we considered that samples presenting reads for a viral family with a read count of less than 1% than the highest count among all libraries sequenced in the same MiSeq run to be product of contamination and hence excluded from further analyses.

Given that *Anelloviridae* was the major viral family identified across the samples (see Results section 4.1), we were prompted to further explore the structure and diversity of the novel viral genomes. The genome sequences were annotated by the Transfer annotations tool in Geneious Prime v.2021.2.2^48^ with a 40% similarity threshold, using as template TTV tamarin (NCBI accession number NC_014085.1) from *S. oedipus*, as well as identifying ORFs through the Find Orf tool in Geneious. Complete genomes were circularized also in Geneious by identifying the GC-rich region in *Anelloviridae*.

### 3.4. Phylogenetic analysis of novel *Anelloviridae* genomes

To contextualize the novel genomes within the Anelloviridae diversity, all annotated contigs were searched with the getorf/EMBOSS v.6.6.0.0^49^ for the presence of ORF1 fragments with at least 1,500 nucleotides. These sequences were aligned to a dataset previously assembled containing all species from the viral family^19^, composing a comprehensive dataset of 618 ORF1 sequences. These sequences were translated and aligned with MUSCLE v.3.8.425^50^. The alignment was trimmed with TrimAl v.1.4^51^ with the ‘--gappyout’ option. A maximum likelihood tree was then inferred with IQ-Tree v.2.1.4 ^52^ under the LG+F+G4 model ^53^, suggested by ModelFinder^54^, and with the implementation of the SH-like approximate likelihood ratio test (SH-aLRT)^55^. A complete list of used sequences, as well as the described alignment and tree are available in Supplementary File S1.

### 3.5. Molecular clock analysis of TTVs from NP

A separate nucleotide dataset comprising all TTVs from NP was assembled and a maximum likelihood tree was inferred under the GTR+F+I+G4 model ^56,57^. The tree and alignment were used for identity analysis with the R software^58^, using the APE package v.5.5 ^59^. As the phylogenetic pattern observed at the genus level was consistent with the ancient within-species diversity model, we were prompted to infer a time-scaled phylogenetic tree for *Episolontorquevirus* genus, which comprehends TTVs from capuchin (*S. nigritus*), tamarin (*S. oedipus*) and the new marmoset (*C. penicillata*) viruses. This analysis was performed with BEAST v1.10.4^60^. Following Perelman et al. (2011)^61^, estimated host divergence dates were used to calibrate internal nodes of the viral tree. A calibration time point between TTV capuchin and other *Epsilontorquevirus* was set by a normal distribution of mean 19.95 million years ago (mya; 95% CI = 15.66 - 24.03 mya), while the calibration time point between TTVs tamarin and marmoset was set by a normal distribution of mean 14.89 mya (95% CI = 11.04 - 18.48 mya). Marginal likelihoods for eight different combinations of tree priors and clock models have been evaluated with path-sampling and stepping-stone sampling algorithms^62,63^. The evaluated tree priors were: speciation yule and coalescent constant, exponential and skyline^64–66^. Both the strict and uncorrelated relaxed clock models have been assessed ^67,68^. For each model combination, a Markov chain Monte Carlo run with 50 million generations was executed, sampling every 5,000 steps. Mixing (effective sample size > 200) for all models parameters was verified on Tracer v.1.7.0^69^ and a maximum clade credibility tree was inferred with TreeAnnotator v1.10.4^69^ for the best fit model. All BEAST xmls, logs and maximum clade credibility are available in Supplementary File S2.

## 4. Results

### 4.1. High-throughput sequencing and taxonomic assignments

The HTS protocol generated a total of 8,352,114 reads, harboring a total of 1,487,358 raw reads for P1 library, 3,319,936 for P2 and 3,544,820 for P7. After reads were filtered, a total of 363,161 (24.4%), 696,397 (21%) and 544,058 (15.3%) reads were obtained for libraries P1, P2 and P7, respectively. Reads were taxonomically assigned through similarity search against the complete nr database from NCBI. Viral reads corresponded to approximately 0.142% of the total generated data (1,109 viral reads in P1, 9,200 in P2 and 1,547 in P7). The meta-spades software *de novo* assembled more than 54 thousand contigs, 15,519 contigs assembled in P1, 17,500 in P2 and 21,671 in P7. In the assembled contigs, 312 (0,57% of total) were taxonomically assigned as viruses, from which 78 (0.5%), 186 (1%) and 48 (0.2%) belonged to the sequenced libraries P1, P2 and P7, respectively.

Several viral families have been identified among the evaluated libraries, with bacteriophage families being the major viral group assigned, corresponding to 70,5%, 60,8% and 79,57% of viral reads and 62,82%, 22,22% and 72,92% for viral contigs in P1, P2 and P7, respectively. Representative reads and contigs for *Siphoviridae*, *Myoviridae*, *Autographiviridae*, *Podoviridae* and *Herelleviridae* were identified. As for vertebrate viruses, diverse reads have been classified as belonging to the families *Retroviridae*, *Papillomaviridae*, *Anelloviridae* and *Genomoviridae*. Among the vertebrate viral families identified, *Anelloviridae* stands out as the one presenting the vast majority of viral reads and contigs (3,816 reads and 164 contigs in total), being hence the main group explored throughout this study. A complete table with viral taxonomic assignments can be found in Supplementary Table S1.

### 4.2. *Anelloviridae* characterization

The *Anelloviridae* family was identified in all evaluated pools, comprehending 216 assigned reads for library P1, 3,514 for P2 and 86 for P7. All of them had a read count greater than the contamination cut-off previously established (*n* = 35), supporting the bonafide presence of the viral family. A total of 27, 129 and 8 assembled contigs were identified as *Anelloviridae* (Table 2) for P1, P2 and P7, respectively. To the best of our knowledge, this is the first report of this viral family in the black-tuffed marmoset (*C. penicillata*).

**Table 2:** Summary of the number of total reads and contigs identified as Anelloviridae and number of ORF1 regions and complete genomes obtained.

| **Library** | **Filtered Reads** | **Anellovirus Reads** | **Contigs** | **Anellovirus Contigs** | **ORF1** | **Genomes** |
| --- | --- | --- | --- | --- | --- | --- |
| P1 | 363,161 | 216 | 15,519 | 27 | 1 | 0 |
| P2 | 696,397 | 3,514 | 17,500 | 129 | 14 | 2 |
| P7 | 544,058 | 86 | 21,671 | 8 | 0 |  |

To further contextualize the novel sequences within the *Anelloviridae* diversity, all contigs were searched for the presence of ORF1 fragments with at least 1,500 nucleotides. We found a total of one contig with an ORF1-like in P1 and 14 contigs in P2. No contig with ORF1-like was identified in P7. Among those 15 contigs, two of them possessed a flanking GC-rich region and all other anellovirus ORFs could be completely annotated, confirming the presence of two whole novel genomes of ~3kb in P2 library (Figure 1, Tables 2 and 3).

**Figure 1:**
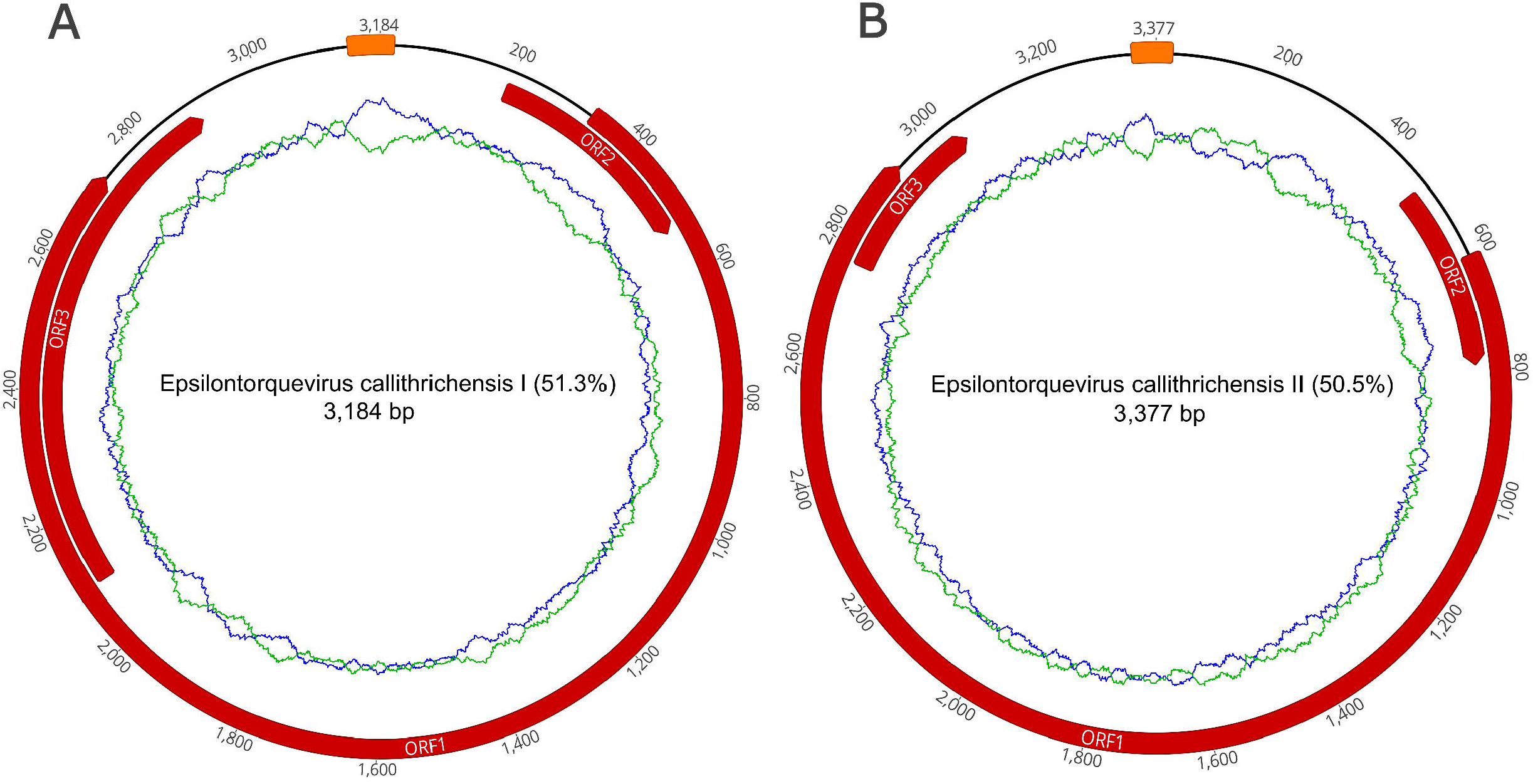
Complete gene maps of the *C. penicillata* torque teno marmoset virus types 1 and 2 (provisionally named *Epsilontorquevirus callithrichensis I* and *Epsilontorquevirus callithrichensis II*) genomes. Gene annotations are represented in red. Both genomes contained three open reading frames: ORF1, ORF2 and ORF3. The putatite non-coding region, Repeat Region, was located between the ORF1 start codon and ORF3 stop codon. The green ring represents the A□+□T content and the blue ring indicates C□+□G content.

**Table 3:** Genome characteristics of the two novel TTV marmosets characterized.

| Type | Genome Length | %GC | Nucleotide position and size<br>(amino acid) |  |  |  | Most closely related<br>Anellovirus | % ORF1 Seq ID |
| --- | --- | --- | --- | --- | --- | --- | --- | --- |
|  |  |  | ORF1 | ORF2 | ORF3 | Repeat<br>Region |  |  |
| *Epsilontorquevirus*<br>*callithrichensis I* | 3.184 bp | 51.3 | 327 - 2.729<br>(801) | 196 - 534<br>(113) | 2.095 - 2.895<br>(267) | 3.137 - 21<br>(69)<sup>a</sup> | TTV Tamarin | 53,49<sup>b</sup> |
| *Epsilontorquevirus*<br>*callithrichensis II.* | 3.377 bp | 50.5 | 615 - 2.927<br>(771) | 484 - 789<br>(102) | 2.758 - 3.039<br>(94) | 3.339 - 26<br>(65)<sup>a</sup> | TTV Tamarin | 51,7<sup>b</sup> |

(a) Nucleotide size of Repeat region
(b) Results are based on ORF1 pairwise identity in Clustal Omega. TTV Tamarin, Torque teno tamarin virus (Accession number NC\_014085)

The two novel genomes sequenced were submitted to ICTV and were provisionally named as *Epsilontorquevirus callithrichensis* (*E. callithrichensis*) types *I* and *II*. The *E. callithrichensis* I is 3,184bp long with a guanine-cytosine (GC) content of 51.3%, while *E. callithrichensis* II is 3,337bp long with a GC content of 50.5%. Both genomes contain the three major ORFs found in *Anelloviridae*. The complete ORF1 gene, a fundamental gene to species determination in the family, had an estimated length of 2,403 and 2,313bp for the *E. callithrichensis* I and II, respectively, and shared less than 53.49% identity to previously known *Epsilontorquevirus*. Particularly, the ORF3 gene from *E. callithrichensis* I had an estimated length of 801bp, superior to the estimated ORF3 in *E. callithrichensis* II with 282bp and to the TTV tamarin (Accession Number NC_014085.1) ORF3, with 497bp.

A viable ORF4 was not found in any *E. callithrichensis*, but the Transfer annotations tool in Geneious software found regions with moderate similarity (52.89%, nucleotide positions 2,189 - 2,896) to TTV capuchin ORF4 in *E. callithrichensis* I. The same region similarity was also found in *E. callithrichensis* II (49.26%, nucleotide positions 2,356 - 3,040), as well as regions with moderate similarity to TTV tamarin ORF4 (48.15%, nucleotide positions: 2,978-3,145). Those hypothetical ORF4 found in *E. callithrichensis* do not possess the typical start codons from other annotated ORFs in *Anelloviridae*.

### 4.3. Evolutionary Analysis

To contextualize the novel *Anelloviridae* genome sequences, a maximum likelihood phylogeny was reconstructed with a comprehensive dataset, following Varsani et al. (2021). The obtained tree presented phylogenetic consistency with the previously proposed *Anelloviridae* phylogeny, being all genera inferred as monophyletic with moderate to high support values (SH-aLRT = 67.4 - 100). The marmoset anelloviruses composed a monophyletic group within *Epsilontorquevirus* (SH-aLRT = 100), a genus comprehending virus species described for the NP. The external group of *Epsilontorquevirus* was the *Zetatorquevirus* genus, which comprehends the TTV douroucouli sampled in *Aotus trivirgatus*^27^ (Figure 2). This major clade comprised all characterized NP TTVs.

**Figure 2:**
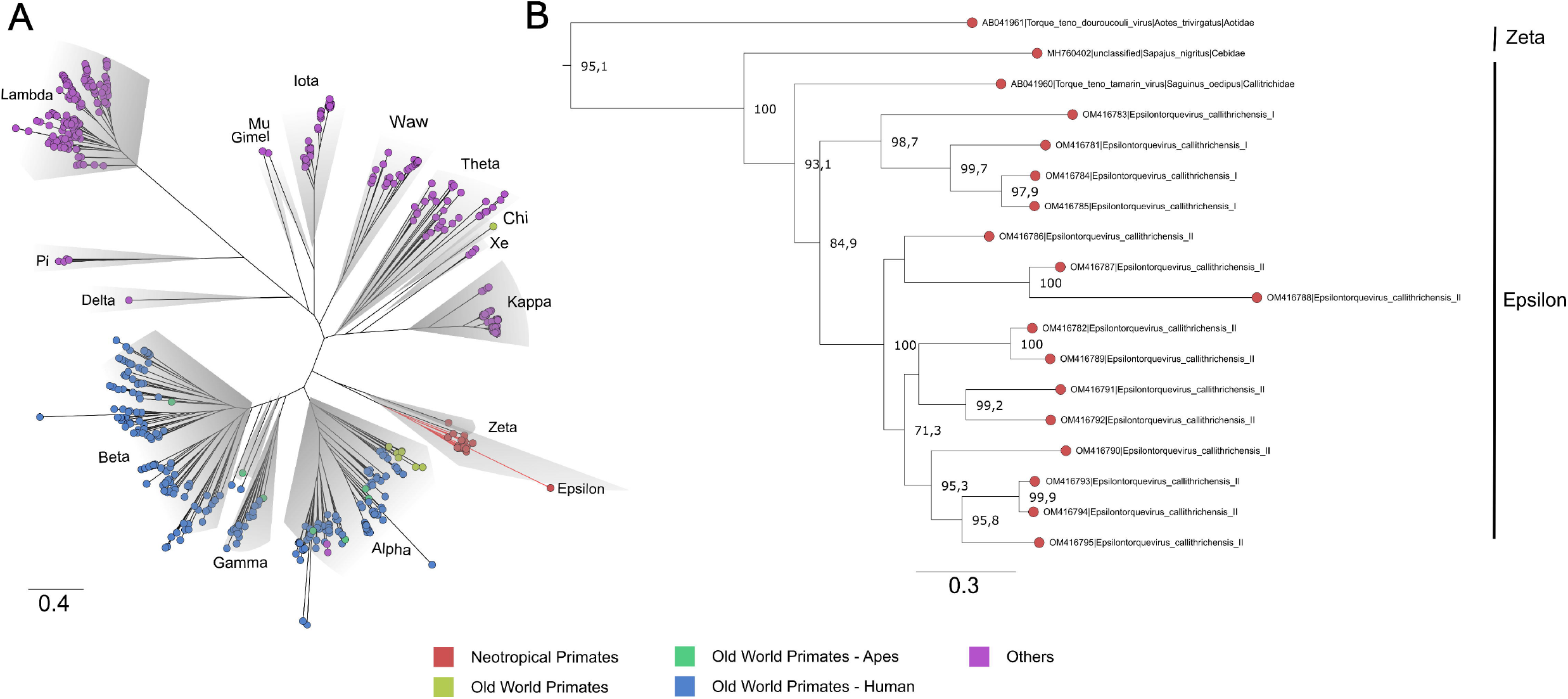
Phylogenetic tree based on complete ORF1 dataset of the *Anelloviridae* family. (A) The *Anelloviridae* tips were colored according to the virus host: strains infecting NP have tips highlighted in red, OWP in yellow, OWP from the Ape lineage in green and Humans in blue. The main representative genera are highlighted in gray. The novel viruses herein described are provisionally named *Epsilontorquevirus callithrichensis I* and *Epsilontorquevirus callithrichensis II* and formed a monophyletic group within *Epsilontorquevirus*. Branches in red are the sequences obtained in this work. (B) The *Zetatorquevirus* genus is the external group from *Epsilontorquevirus*, which is divided in three clades according to host species, with great genetic divergence in the *Epsilontorquevirus callithrichensis* lineage.

To verify whether the newly identified marmoset viruses belonged to a new species or genus, a separate dataset for NP TTVs was assembled and analyzed. Pairwise nucleotide sequence distances were obtained and visualized through heatmap (Figure 3). The ORF1 sequences from the novel viruses clustered in two monophyletic groups, presenting pairwise identities between 55.89 and 59.35% to the previously described *Epsilontorquevirus* in tamarin. For the TTV capuchin, distances ranged from 53.68 to 56.58%. The ICTV criteria for TTV classification establishes genetic distances thresholds for species (69%)^19^. When compared to the single *Zetatorquevirus* described, the marmoset anelloviruses demonstrated diminished identity, ranging from 42,19% to 44,96%. The first marmoset anellovirus monophyletic lineage, named *E. callithrichensis I*, had a pairwise nucleotide sequence distance ranging from 65.68% to 86.92% among its representatives. The second, named *E. callithrichensis II*, had a pairwise nucleotide sequence distance ranging from 60.66% to 88.89% among its representatives. In this sense, the results presented in this study indicate the existence of two novel viral species from the genus *Episolontorquevirus* - with a 57.28% to 61.3% pairwise nucleotide distance. Additionally, we report considerable intra-species genetic differences, suggesting that diverse viral strains co-circulate. The pairwise identity matrix of NP *Anellovirus* can be found in the Supplementary File S1.

**Figure 3:**
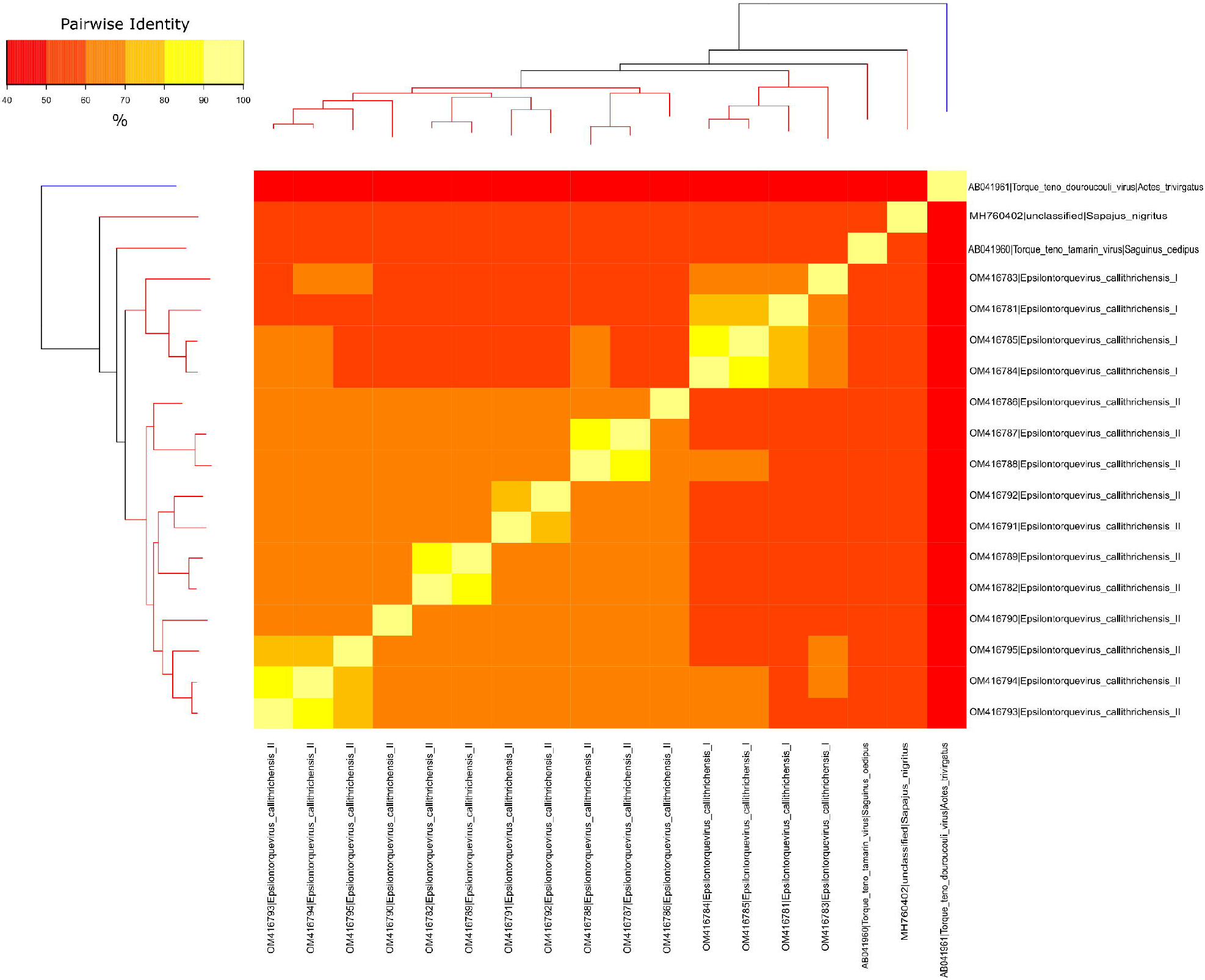
Phylogenetic tree and heatmap displaying the pairwise nucleotide sequence distances of NP anelloviruses. The phylogeny represented both on the left and upper corners is a Maximum likelihood tree of the known anelloviruses in NP. Branches in red are classified as *Epsilontorquevirus*, while branches in blue are classified as *Zetatorquevirus*. Sequence distances are indicated by different colors in the central plot. The *Epsilontorquevirus* was composed of four main lineages, one previously discovered in *Sapajus nigritus* and one in *Saguinus oedipus*, with the two newly discovered in *C. penicillata* in this work. A great genetic diversity may be observed among *C. penicillata* anelloviruses, such that they can be classified as belonging to two new species, named: *Epsilontorquevirus callithrichensis I* and *Epsilontorquevirus callithrichensis II*.

### 4.4. Timescale Analyses

The phylogenetic pattern inferred for the genus *Epsilontorquevirus* (Figure 2B) was consistent with the topology of their host species tree, echoing the within-species diversity followed by co-divergence model of viral diversification. The TTV capuchin was the first lineage to diverge within *Epsilontorquevirus*, followed by TTV tamarin, which is sister to the clade comprehending all TTVs marmoset. This branching pattern is consistent with the Cebidae molecular phylogeny inferred^61^. In this sense, we inferred a time-scaled phylogenetic tree for *Episolontorquevirus* genus using host divergence dates to calibrate internal nodes of the viral tree. Among all models tested, the one with the best statistical fit was characterized by the strict clock and the Yule speciation prior (Supplementary file S2, Path and Stone Sampling table).

Overall, our time scale inference found that the most recent common ancestor of the *Episolontorquevirus* genus was estimated around 18.7 mya (95% HPD = 15.31 - 21.88), in accordance with the prior established for the analysis. In addition, the divergence across the TTV tamarin and *E. callithrichensis* was estimated around 15.48 mya (95% HPD = 12.73 - 18.17). The divergence between the *E. callithrichensis I* and *E. callithrichensis II* was estimated around 13.1 mya (95% HPD = 10.73 - 15.50 mya). The ancestor for *E. callithrichensis I* was dated around 7.9 mya (95% HPD = 6.37 - 9.54 mya) and the one for *E. callithrichensis II* around 9.6 mya (95% HPD = 7.71 - 11.2 mya) (Figure 4).

**Figure 4:**
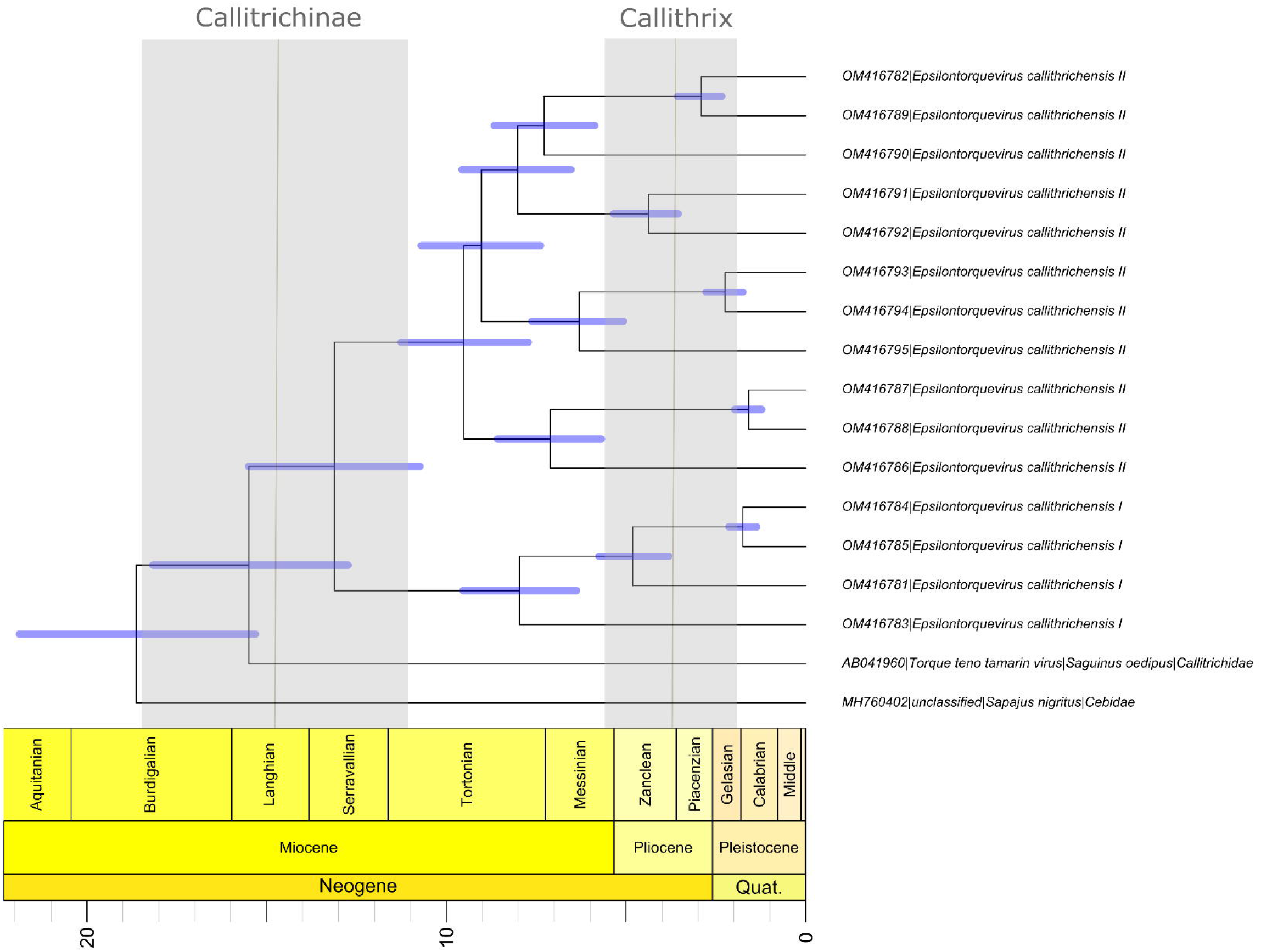
Divergence time estimates for *Epsilontorquevirus* based on host divergence times used for calibrating the viral molecular clock. Branch lengths are proportional to divergence times and blue bars represent the 95% HPD interval of virus divergence times. The black line represents the divergence times of Callithrix and Callitrichidae, while the blue columns represent its 95% HPD interval, as proposed by Perelman et al., 2011.

## 5. Discussion

With the development of HTS protocols to explore the diversity of existing viruses, previous works have identified new anelloviruses species in wildlife - bats, rodents^38^, seals^29^, wild felids^30^, canids^31^ and non-human primates ^9,10,25,27,32,33^. In this work, we sequenced and characterized two novel *Anelloviridae* species genomes in NP, increasing the host range for this family. Although the diversity of TTVs in primates has been previously explored, especially for OWP ^25–27^, the known diversity in NP is yet remarkably limited. Until the present study, only two studies identified anelloviruses in NP ^10,27^, though no effort to characterize their evolutionary dynamics has been presented. Hence, this is the first attempt to investigate in depth the evolution of novel NP anellovirus.

The anelloviruses sequenced in this work were unique to the known NP *Anelloviridae* diversity, comprehending two novel species by the ICTV species criterion (Figure 1 and 2, Table 3). Both viruses’ complete genomes had the expected ORFs found in this viral family: the putative capsid protein and replication-associated protein (ORF1) and the ORF2 protein^70^, as well as the putative ORF3. By particularly looking at some regions of the novel genomes, we could identify regions similar to TTV tamarin and TTV capuchin ORF4, though with numerous stop codons and frameshift mutations, implying lack of transcriptional function. This suggests ORF4 is not fundamental for *E. callithrichensis* infection and shows an instance of genomic structure plasticity in ssDNA viruses. The adoption of models and cell systems to study anelloviruses infections in vitro shall improve the knowledge regarding the function and structure of its proteins, deeply scarce at the present time.

Phylogenetic analysis of the new genome sequences suggests a topological pattern consistent with a previously proposed model for the diversification of multiple viral families, the within-species diversity followed by codivergence model ^16,40–42^. Under this model, viral lineages diversify either alongside their host species or through a speciation process within a single host, occasionally caused by niche adaptation to specific cell types or tissues. This process culminates in a viral phylogenetic pattern where multiple clades reflect the host phylogeny, as is the case of primate TTVs^27^. We explored the consequences of this model to calibrate the molecular clock and obtain, for the first time, a timescaled phylogenetic tree for NP TTVs (Figure 4). This analysis suggests that the lineages that lead to both species described in this study diverged around 13.1 mya, while the time for the ancestors of *E. callithrichensis I* and *II* were dated around 7.9 and 9.6 mya, respectively. Both dates are much older than the origins of *Callithrix*, implying that these viral lineages circulated among the ancestors of *Callitrichinae* (around 14.9 mya)^61^. We thus hypothesize that its descendant extant species may harbor TTVs related to the described viral lineages (Figure 5).

**Figure 5:**
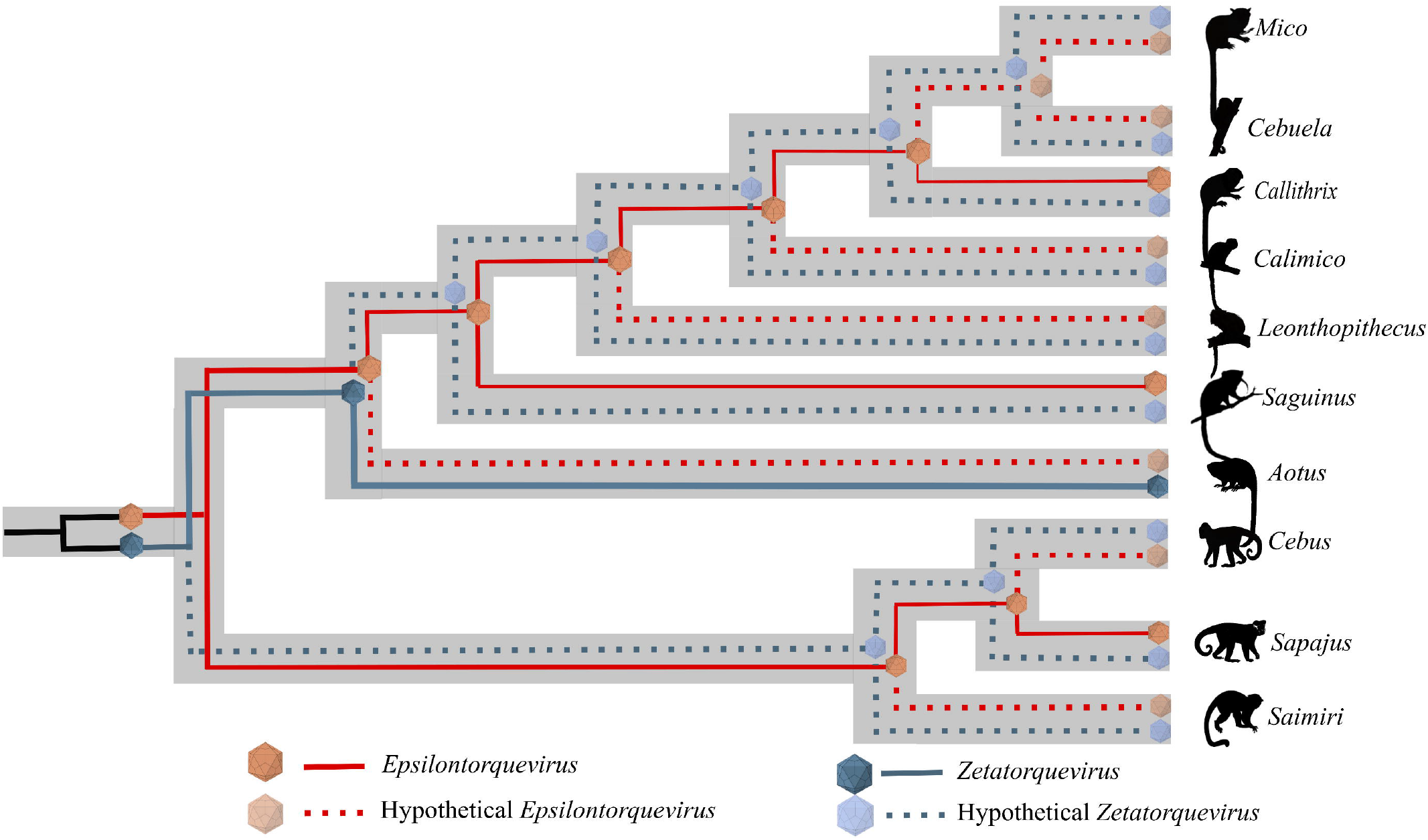
Schematic model of ancient within-species diversity model followed by codivergence in NP from the *Cebidae* family. Continuous lines represent strains already sampled and dashed lines represent putative strains that might explain more likely ancient ancestors. The *Epsilontorquevirus* genus is represented by red lines, *Zetatorquevirus* by blue lines. We hypothesize the existence of an ancestral *Anelloviridae* in the Cebidae ancestor, that posteriorly originated *Epsilontorquevirus* and *Zetatorquevirus* in a mirrored evolutionary process.

We observed a high genetic diversity between the *E. callithrichensis* characterized herein (Figure 3), a phenomenon already described in anelloviruses due to their high mutation rate ^71–73^. The presence of multiple lineages co-infecting a host has already been observed as three distinct *Anelloviridae* genera were described in humans ^74^. The same pattern is present in OWP, as chimpanzees are infected by *Alpha, Beta* and *Gammatorquevirus* ^24–27^. Once the NP *Anelloviridae* diversity is further explored, and the diversity of *Epsilontorquevirus* and *Zetatorquevirus* is thoroughly addressed, the pattern of coinfection by multiple TTV strains may be further verified in NP (Figure 5).

While previous observations of codivergence patterns have been made for other TTVs^27,30^, no clear evolutionary model has been proposed to assess TTV diversity. Although this study focused on NP TTVs, we argue that the within-species diversity followed by codivergence model might explain the drivers of diversification for the whole *Anelloviridae* family. Further characterization of TTVs diversity in non-model organisms through metagenomics might play a key role in unraveling the validity of the proposed model. Finally, this study showcases an instance of the importance of sampling wildlife viruses to obtain a better characterization of the long-term evolutionary processes driving the diversification of viral families.

## 7. Data Availability

Sequences generated in the present work were submitted to Genbank under the accession numbers: OM416781-OM416795. Supplementary files can be found in https://github.com/matheus-cosentino/Cosentino_2022_Epsilontorquevirus.

## 8. Acknowledgements

This work was supported in part by the Conselho Nacional de Desenvolvimento Científico e Tecnológico/CNPq (grant 313005/2020-6 to A.F.A.S.) and Fundação de Amparo à Pesquisa do Estado do Rio de Janeiro/FAPERJ (grants E-26/112.647/2012 to M.A.S. and E26/202.738/2018, E26/210.122/2018 and E26/211.040/2019 to A.F.A.S.) for funding this study. We thank to all the workers of the Centro de Primatologia de Brasília, Universidade de Brasília (Distrito Federal, Brazil) for assistance with sample collection and to the fantastic LDDV members, in special to Déa Mota, for the wonderful team work. We are also grateful to Carolina Furtado and Nicole Scherer from Instituto Nacional de Câncer Jośe de Alencar (Rio de Janeiro, Brazil) for assistance with HTS and analysis.

